# Cardiomyocyte differentiation from iPS cells is delayed following knockout of Bcl-2

**DOI:** 10.1101/2022.01.05.475068

**Authors:** Tim Vervliet, Robin Duelen, Rita La Rovere, H. Llewelyn Roderick, Maurilio Sampaolesi

**Affiliations:** Laboratory of Molecular and Cellular Signaling, Department of Cellular and Molecular Medicine, KU Leuven, 3000 Leuven, Belgium; Translational Cardiomyology Laboratory, Department of Development and Regeneration, KU Leuven, 3000 Leuven, Belgium; Laboratory of Experimental Cardiology, Department of Cardiovascular Sciences, KU Leuven, 3000 Leuven, Belgium

## Abstract

Anti-apoptotic B-cell lymphoma 2 (Bcl-2) regulates a wide array of cellular functions involved in cell death, cell survival decisions and autophagy. Bcl-2 acts by both direct interaction with different components of the pathways involved and by intervening in intracellular Ca^2+^ signalling. The function of Bcl-2 is in turn regulated by post-translational modifications including phosphorylation at different sites by various protein kinases. Besides functions in cell death and apoptosis, Bcl-2 regulates cell differentiation processes, including of cardiomyocytes, although its mechanism of action in this process is not fully elucidated. To further address the role of Bcl-2 in cardiomyocyte differentiation, we investigated the effect of its genetic knockout by CRISPR/Cas9 on the differentiation and functional maturation trajectory of human induced pluripotent stem cells (hiPSC) to cardiomyocytes. Our results indicate that differentiation of hiPSC to cardiomyocytes is delayed by Bcl-2 KO. This effect was associated with smaller areas of spontaneously beating cells. At the cell population level this exhibited in reduced expression and activity of the cardiomyocyte Ca^2+^ toolkit. However, when restricting the comparison to the active areas exclusively Bcl-2 KO did not show to significantly alter the functionality of the differentiated cardiomyocytes. Finally, Bcl-2 KO reduced c-Myc expression early in the cardiac differentiation process which may account at least in part for the here observed delay in cardiac differentiation resulting reduced efficiency of the differentiation process. These data are supportive of a pivotal role for Bcl-2 in cardiomyocyte differentiation and maturation.

## Introduction

B-cell lymphoma 2 (Bcl-2) is involved in the regulation of cell death and survival decisions at multiple levels. Canonically, the anti-apoptotic Bcl-2 protein binds to pro-apoptotic Bcl-2-family members, such as Bax and Bak, thereby inhibiting their pro-apoptotic functions as inducers of mitochondrial outer membrane permeabilization (MOMP) and thus cytochrome C release (1). In addition, Bcl-2 modulates inositol-1,4,5-trisphosphate receptors (IP_3_R) thus, inhibiting pro-apoptotic Ca^2+^ signalling towards the mitochondria (2). Other Ca^2+^ handling proteins targeted by Bcl-2 include ryanodine receptors (RyR), sarco/endoplasmic reticulum ATPases and voltage dependent anion channels (3). Besides regulating the apoptotic outcome, Bcl-2 is involved in modulating autophagic flux. For instance, Bcl-2 binds to Beclin 1, thereby limiting the initiation of autophagosome formation and thus autophagic flux (4).

Regulation of cell death and survival is also key during differentiation. Consistently, Bcl-2 is involved in the differentiation of certain cells and tissues. Neuronal differentiation for instance, has been shown to be regulated by Bcl-2. Transplanting embryonic stem cells overexpressing Bcl-2 in rat cortex enhanced recovery and neuronal differentiation after ischemic insult (5). In addition, overexpression of Bcl-2 in the PC12 neural crest tumour cell line, led to increased expression of genes associated with neural differentiation, whereas it decreased proliferation-related genes (6). In the heart, Bcl-2 was shown to be upregulated in a GATA4-dependent manner (7), suggesting a role for it in cardiac development. The latter is further supported by the increased cardiomyocyte proliferation seen in mice overexpressing Bcl-2 in the heart (8). However, the molecular pathways underlying Bcl-2 regulation of cardiomyocyte differentiation remain elusive.

Bcl-2 is phosphorylated at a number of sites resulting in modulation of its function. For example, Thr56, Ser70 and Ser87 located in the Bcl-2 unstructured loop are targets of multiple kinases including members of the mitogen-activated protein kinases (MAPK) family of kinases (9-11). Phosphorylation of Bcl-2 by p38MAPK for example was demonstrated to induce apoptosis by inhibiting the anti-apoptotic function of Bcl-2 and inducing it to trigger the release of cytochrome C following MOMP induction in response to acute stressors (11). Phosphorylation of Bcl-2 has also been shown to result in its translocation to the nucleus (12). In the nucleus, Bcl-2 interacts with c-Myc increasing its stability and transcriptional activity (13). These functional interactions between Bcl-2, p38MAPK and c-Myc likely also play key role in the heart. Indeed, p38MAPK commits mouse embryonic stem cells to differentiate into cardiomyocytes (14) and regulates cardiomyocyte proliferation (15). Whereas, cardiac c-Myc overexpression in mice increased cardiac mass and hyperplasia without detrimental effects on cardiomyocyte maturation (16).

In this study, we aim to further unravel the function of Bcl-2 in cardiomyocyte differentiation. To this end, we analysed the effect of CRISPR/Cas9-mediated Bcl-2 knock-out (KO) on the derivation of cardiomyocytes by differentiation from human induced pluripotent stem cells (hiPSC). We show that the temporal trajectory of hiPSC differentiation towards cardiomyocytes is delayed by Bcl-2 KO, resulting in functional changes. We further show that Bcl-2 KO cells have reduced c-Myc expression during early cardiomyocyte differentiation compared to control cells. Moreover, in control cells this increased c-Myc expression coincides with upregulation of Bcl-2 levels. Together, these results support a critical role for Bcl-2 in the generation of cardiomyocyte differentiation from hiPS and thus suggest a role in differentiation of cardiomyocytes during embryonic development.

## Materials and methods

### Reagents and antibodies

Unless otherwise specified, all chemicals were purchased from Merck. All primers and gene blocks used in this study were purchased from integrated DNA technologies (IDT; Leuven, Belgium). Primers were listed in (Supplemental table S1 and S2). Primary antibodies were anti-Vinculin (#V-9131), anti-c-Myc (#M4439) (Merck), anti-Tubulin (BD Pharmingen; #556321) anti-Bcl-2 (#sc7382HRP), anti-phospho Ser70-Bcl-2 (#sc-293128) (Santa Cruz), anti-phospho-p70 S6 kinase (#9234S), anti-p70 S6 kinase (#9202), anti-phospho-p38MAPK (#9215S), anti-p38MAPK (#9212), anti-Bcl-X_L_ (#2764), anti-PARP (#9532S) (Cell Signalling), anti-LC3 (Nanotools; #0231-100/LC3-5F10), Anti-RyR (C3:33) (#MA3-916), anti-NCX (#C2C12) (ThermoFisher), anti-cTnT (Abcam; #ab209813), anti-IP_3_R2 (Abiocode; # R2872-2) and anti-SERCA2a (produced and kindly gifted by Prof. F. Wuytack (KU Leuven)).

### hiPSC culturing

All experiments were performed using the commercially available Gibco™ Episomal hiPSC Line (A18945; Thermo Fisher Scientific). hiPSC were cultured feeder-free on Geltrex LDEV-Free hESC-Qualified Reduced Growth Factor Basement Membrane Matrix (ThermoFisher; #A1413201) and maintained in Essential 8 Flex Basal Medium supplemented with Essential 8 Flex Supplement (50x) and 0.1% Pen/Strep (all from Thermo Fisher Scientific), at 37°C under normoxic conditions (21% O_2_ and 5% CO_2_). Medium was changed daily. Colonies were routinely passaged non-enzymatically with 0.5 mM EDTA in Phosphate-Buffered Saline (PBS) (Thermo Fisher Scientific).

### Cardiomyocyte differentiation of hiPSC

For inducing cardiac differentiation, the iPSC cardiomyocyte differentiation kit (Thermo Fisher Scientific; #A2921201) was used according to the manufacturers protocol. Briefly, prior to differentiation, hiPSC cells were seeded on a thin Matrigel™ Growth Factor Reduced (GFR) Basement Membrane Matrix layer (Merck; #CLS356252) and cultured for 3 or 4 days in Essential 8 medium at 37°C under hypoxic conditions (5% O_2_ and 5% CO_2_) until the cells reached approximately 60% of confluency. Cardiomyocyte differentiation was started after the addition of a mesoderm-inducing medium (medium A; day 0) for 48 h. After 24 h, cells were transferred from hypoxia to normoxia. At day 2 of differentiation, cells were incubated for another 48 h with a cardiomyocyte progenitor medium (medium B). From day 4 onwards, medium was changed every other day with a cardiomyocyte maintenance medium. For experimental purposes, samples were harvested via EDTA incubation and scraping on different days (0, 2, 4, 7, 14 and/or 21). All experiments performed were approved by the Research Ethical Committee of UZ/KU Leuven (protocol number S62524).

### Cloning of gRNA and selection plasmid

The Bcl-2 gRNA (forward: 5’-GAGAACAGGGTACGATAACC-3’ and reverse: 5’-GGTTATCGTACCCtGTTCTC-3’) was cloned in the pU6-(BbsI)CBh-Cas9-T2A-mCherry, a gift from Ralf Kuehn (Addgene plasmid # 64324 ; http://n2t.net/addgene:64324 ; RRID:Addgene_64324) (17), using the BbsI restriction enzyme. In order to introduce a hygromycin selection cassette into the genomic *BCL2* sequence, two genomic DNA sequences, in close proximity and on either side of a TTAA sequence within 500 bp of the Bcl-2 gRNA, were identified. These sequences (one 430 bp (HAL sequence (see ‘Supplemental materials and methods’) and a 450 bp long (HAR sequence)) were introduced using the NEBuilder^®^ HiFi DNA Assembly Master Mix (New England BioLabs) in combination with the sequences produced as gene blocks (IDT) and the PiggyBAC-Hygro-TK vector flanking the hygromycin resistance gene restricted with BamHI enzyme. In the HAR sequence, containing the start of the *BCL2* open reading frame, base pair substitution were incorporated in order to generate stop codons early in the *BCL2* gene resulting in the knock-out.

### CRISPR/Cas9-mediated knock out of Bcl-2

Transfection of hiPSC was performed by nucleofection using the commercially available Cell Line Nucleofector Kit (Lonza) according to their protocol. For the nucleofection, 2 µg of the gRNA containing plasmid and 8 µg of the selection plasmid were utilized. After nucleofection, the hiPSC cells were plated onto Geltrex coated culture plates. On the third day after nucleofection, 50 ng/ml hygromycin was added to the medium to start the selection procedure. After 7 days of selection, single colonies were picked. The medium was refreshed each day with Essential 8 medium containing 50 ng/ml hygromycin until the colonies were sufficiently grown, after which cells were passaged. At this point, a fraction of the hiPSC was taken for genomic DNA isolation to screen for the insertion of the hygromycin selection cassette at the correct locus in the *BCL2* gene (see Supplemental table S1). Finally, in clones selected based on correct insertion of the hygromycin gene in the *BCL2* locus, the selection cassette was again removed. To this end, hiPSC cells were detached and nucleofected with 5 µg pf the piggyback transposase plasmid (kind gift of Dr. Catherine Verfaillie (KU Leuven)). The medium was changed daily using Essential 8 medium until cells reached 90% confluency. At this point Fialuridine (FIAU; Merck) selection was started for 24 h to kill cells in which the selection plasmid was retained. After this, the cells were allowed to recover and grow with daily medium changes followed by single colony picking. Finally, a complete gene editing-free Bcl-2 KO hiPSC line was obtained due to PiggyBac excision and FIAU selection. Once the colonies were amplified, genomic DNA was collected in order to screen for the absence of the selection cassette, modification in the *BCL2* gene and potential off target genomic insertions/modifications (see Supplemental table 1).

### Genomic DNA isolation

Genomic DNA was isolated using a PureLink™ Genomic DNA Mini Kit (Invitrogen) according to the manufacturers protocol. The primers utilized for the subsequent PCR reactions can be found in Supplemental table S1. Sequencing of the PCR samples was performed by LGC genomics.

### Immunoblot analysis

Cell pellets collected at 0, 7, 14 and 21 days of differentiation were resuspended on ice in lysis buffer (20 mM Tris HCl, pH7.5, 150 mM NaCl, 1.5 mM MgCl_2_, 0.5 mM DTT, 1% Triton X-100 containing both protease (EDTA free protease inhibitors, Thermo Fisher Scientific) and phosphatase (PhosSTOP, Roche) inhibitors), followed by further homogenizing using an eppendorf douncer in order to facilitate lysis. Lysis was performed for at least 30 min with head over end mixing at 4°C followed by centrifugation for 5 min at 4°C at >5000 xg. The supernatant was collected and protein concentration determined using a standard Bradford assay. Immunoblot samples were prepared at 0.5 µg/µl and were resolved by SDS-PAGE on 4-12% Bis Tris gradient gels (ThermoFisher) and transferred to 0.45 µm PVDF (Merck) as described in (18). Following this primary antibody staining was performed over night at 4°C and secondary antibody staining with HRP conjugated antibodies (Bioke) was performed at room temperature for at least 2 h. Immunoreactive bands were visualized using Pierce™ ECL Western Blotting Substrate (Thermo Fisher Scientific) and imaged using a ChemiDoc™ MP imaging system (BioRad).

### Quantitative reverse transcription PCR (RT-qPCR)

RNA isolation was performed using a GenElute™ mammalian total RNA miniprep kit (Sigma) according to the manufacturers protocol. After this, a DNA-free™ kit (Invitrogen) was used according to the manufacturers protocols to remove potential contaminating genomic DNA. RNA concentrations were determined using a Nanodrop. cDNA was prepared by reverse transcription from 1000 ng total mRNA using the High Capacity cDNA reverse transcription kit (Applied Biosystems). For qPCR, forward and reverse primers for the genes of interest (see Supplemental table S2) were mixed with FastStart Universal SYBR Green Master (Rox; Roche). 5 µl of this mixture was pipetted in duplicate per condition in a 384 well plate after which which 5 µl of a 1:10 dilution of the prepared cDNAs were added. Reactions were performed using a ViiA7 Real-Time PCR System (Thermo Fisher Scientific). For analysis, ΔΔCT values were determined for each condition using *GAPDH* and *RPL13a* as reference genes.

### Immunofluorescence staining

Control and Bcl-2 KO cardiomyocytes were stained for cTNT after 14 days of differentiation. Briefly, the cells were fixed with 4% paraformaldehyde for 10 min followed by permeabilization with 0.2% 0.2% Triton X-100 in PBS containing 1% bovine serum albumin (BSA) for 15 min. Blocking of non-specific protein binding sites was performed for 1 h in 10% goat serum. Next, the cells were incubated overnight at 4°C with cTNT antibody followed by the appropriate Alexa-555 conjugated secondary antibody (Thermo Fisher Scientific; 4 *µ*g/mL) for 1 h at 37°C. Confocal images were taken using a a 63x 1.4 NA oil immersion objective on a Zeiss LSM510 confocal microscope.

### Intracellular Ca^2+^ imaging

Intracellular Ca^2+^ measurements were performed in hiPSC-derived cardiomyocytes differentiated for 9 or 11 days. In these experiments, cells were loaded with 1 µM of the Ca^2+^ reporter Fluo-4 AM (solubilized in cardiomyocyte maintenance medium) for 45 min in a humidified incubator at 37°C and 5% CO_2_. Next, the cells were washed twice with cardiomyocyte maintenance medium, after which the dye was de-esterified for 45 min under the same conditions as for loading. Immediately prior to starting the Ca^2+^ imaging experiments, the maintenance medium was replaced with a pre-warmed (37°C) modified Krebs-Ringer solution (135 mM NaCl, 6.2 mM KCl, 1.2 mM MgCl_2_, 12 mM HEPES, pH 7.3, 11.5 mM glucose and 2 mM CaCl_2_). Additions were performed as indicated in the figure. Tetracaine was solubilized in the above modified Krebs-Ringer solution at 1 mM final concentration. Imaging was performed using a Nikon eclipse Ti2 inverted fluorescence microscope (Nikon) equipped with excitation filter FF01-378/474/554/635 and dichroic mirror FF01-432/515/595/730 and emission filter 515/30, all from Semrock. Excitation was performed at 470 nm using a CoolLed pR-4000 (CoolLed). Acquisition of the emitted fluorescent signal was performed at 10 Hz using a pco.edge 4.2bi sCMOS camera (pCO). For image analysis, FIJI software (19) was utilized. To visualise fluorescence changes, a pseudo-line scan was first generated by re-slicing of the image stack at the location of a straight line that was drawn from top to bottom over the largest area of active hiPSC-derived cardiomyocytes. Profiles of fluorescence intensity were plotted and measured as indicated in the figures. To normalise for baseline fluorescence, traces are shown as F/F0where the F0value was obtained during the quiescent period after tetracaine administration. To determine the area of cells showing spontaneous Ca^2+^ activity, maximum intensity Z projections were performed of the first 300 images per stack. A threshold for response was determined for one control condition in each experiment and applied to all other stacks. After thresholding, particle analyses was performed to determine the regions of interest from which the size of the areas were calculated.

### Statistical analysis

Statistical analysis was performed using GraphPad Prism software (Version 9.1.1). All datasets were tested for normal distribution using Shapiro-Wilk tests. For normally distributed data sets showing insignificant variability between the conditions, ANOVA tests were performed. Non normally distributed data sets were analysed using non-parametric Kruskall Wallis tests. Additonal information on the performed statistical tests can be found in the figure legends. If not otherwise specified, p values < 0.05 were considered significant (with p<0.05, p<0.01, p<0.001 and p<0.0001 are designated as *, **, *** and **** respectively).

## Results

### Creation and validation of Bcl-2 KO hiPSC

Bcl-2 KO was performed in the commercially available Gibco™ Episomal hiPSC Line (A18945) using a CRISPR/Cas9 based approach. After screening for their efficiency in HEK293 cells the most promising Bcl-2 gRNA, targeting an early sequence in the *BCL2* gene, was selected and cloned into a vector co-expressing the Cas9 gene. The Bcl-2 gRNA plasmid was introduced into the hiPSC line together with a plasmid containing a hygromycin selection cassette flanked by DNA regions homologous to the Bcl-2 gene allowing the targeted introduction of the hygromycin cassette into the *BCL2* gene. Point mutations, resulting in early stop codons in the *BCL2* gene were introduced in one of the homologous regions, in order to introduce mutations in the *BCL2* gene. After antibiotic selection and single cell colony isolation, the insertion of the hygromycin cassette at the correct genomic locus was validated. Next, in the validated clones the hygromycin cassette was removed followed by FIAU selection, which kills cells containing the selection cassette. After single colony isolation, clones were screened for alterations in the *BCL2* gene. In figure 1, we restricted the validation to the two clones, Bcl-2 KO1 and Bcl-2 KO2, used in this study. At the genomic level in all the tested clones, removal of the hygromycin cassette resulted in the base pair deletion or frame shift in the *BCL2* gene. The majority of the tested clones showed a deletion of 4 base pairs (Figure 1A) resulting in the early introduction of three stop codons at the protein level (Figure 1B). When verified at the protein level via immunoblot, both clones showed absence of endogenous Bcl-2 (Figure 1C), suggesting a complete knock-out of the *BCL2* gene. Finally, we screened for potential off-target effects of the used gRNA by sequencing the 10 most likely off targets of the gRNA in coding genes (Figure 1D). In both clones no alterations near the potential off target sites were detected, suggesting that the Bcl-2 KO occurred specifically in these cells and that these clones are suitable for use.

**Figure 1:**
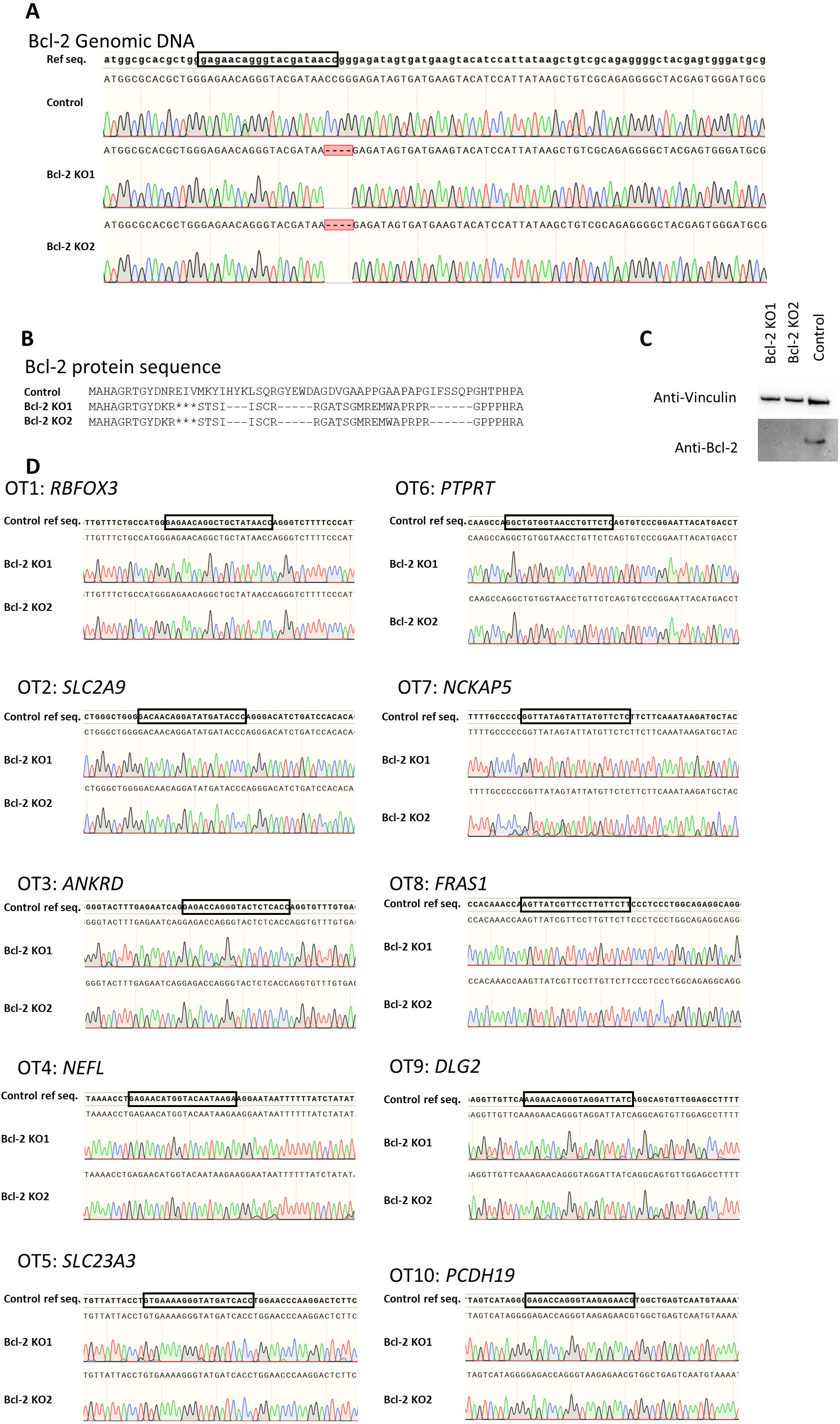
CRISPR/Cas9-mediated KO of Bcl-2 in hiPSC. **A)** Sanger sequencing of the first 100 base pairs of the genomic Bcl-2 open reading frame comparing the sequence of the control condition to the two Bcl-2 KO hiPSC clones utilized in this study. The utilized gRNA is indicated by the black box in the *BCL2* reference sequence (ref seq.). **B)** Predicted protein sequence based on the sequencing result obtained in A for the control and Bcl-2 KO hiPSC. * indicates the presence of a stop codon. **C)** Immunoblot validating the absence of Bcl-2 in the Bcl-2 KO clones compared to the control. **D)** Sequencing results surrounding the 10 most likely off target genes associated with the utilized gRNA. The potential off target is indicated by the black box in the ref seq. All DNA sequences and profiles were visualized using SnapGene^®^ software

### Bcl-2 KO does not inhibit but delays the induction of cardiac differentiation

After validating and the selection of two Bcl-2 KO clones, we set out to determine the effects of Bcl-2 KO on cardiomyocyte differentiation. The differentiation scheme utilized in this study is depicted in Figure 2A. A commercially available cardiomyocyte differentiation kit, which utilizes three different types of medium for differentiation induction, was utilized and resulted in the robust generation of cardiomyocyte cultures.

**Figure 2:**
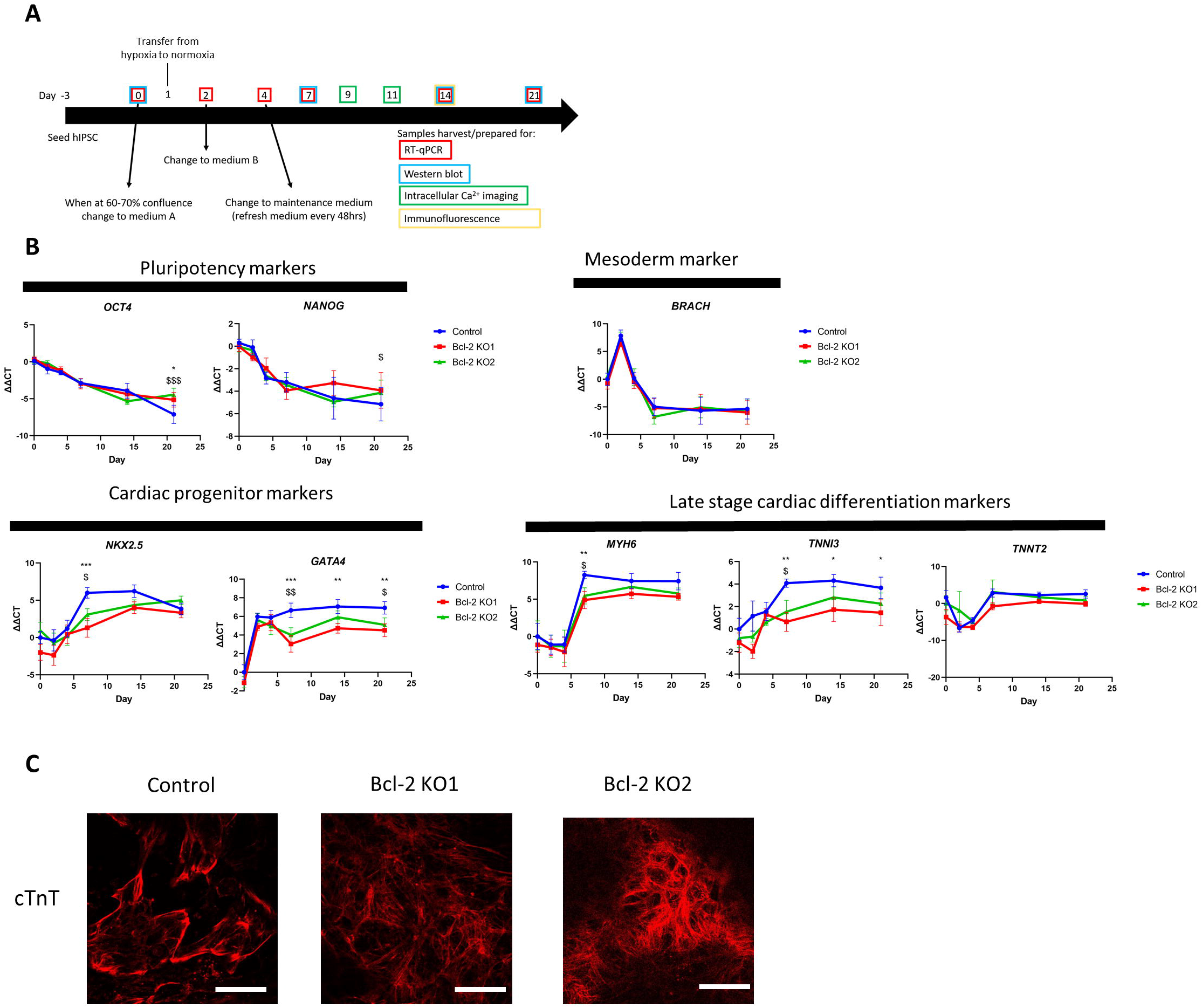
Bcl-2 KO does not impair the expression of cardiac differentiation markers. **A)** Scheme of the utilized cardiac differentiation protocol and time points where samples were harvested for RT-qPCR, immunoblot, immunofluorescence staining or utilized for intracellular Ca^2+^ imaging. **B)** RT-qPCR analysis of gene expression profile associated with cardiomyocyte differentiation in control and the two Bcl-2 KO clones at time points shown (0, 2, 4, 7 14, 21 days of differentiation). The data points represent ΔΔCT ±SEM values of at least 4 different differentiations (n≥4) of the indicated genes at the indicated timepoints. Two way ANOVA tests with Dunnet-post test for multiple comparison were performed * and $ indicate significant difference between control and Bcl-2 KO1 or Bcl-2 KO2 respectively * or $ indicates p<0.05, ** or $$ indicates p<0.01, *** or $$$ indicate p<0.001. **C)** Confocal images of control and Bcl-2 KO cardiomyocytes differentiated for 14 days immunofluorescently stained for cTnT. Experiments were performed 3 times independently (n=3). The white scale bar corresponds to 50 µM

Using this protocol, spontaneous beating of large areas of cardiomyocytes was observed as early as day 7 in the control differentiation, as reported by the manufacturer of the differentiation kit. In the Bcl-2 KO clones, spontaneous beating was not seen until at least 9 days after the start of differentiation. Moreover, the regions of spontaneously active cardiomyocytes were scarcer and smaller (see also figure 4A, B). This would suggest that Bcl-2 KO impact cardiac differentiation. To investigate this further, we first assessed whether Bcl-2 KO resulted in impaired induction of expression of early and late cardiomyocyte differentiation markers (Figure 2B). In control and Bcl-2 KO conditions, RT-qPCR analysis revealed a decrease in expression of pluripotency markers upon differentiation as expected. mRNA expression of the mesodermal marker *BRACH* indicated mesoderm induction occurred as anticipated on day 2 of differentiation for both control and KO conditions. At day 7, early cardiac differentiation markers such as *NKX2*.*5* and *GATA4* are upregulated and should be expressed throughout the remainder of the differentiation. This was indeed the case for the control conditions. However, in both Bcl-2 KO lines, *NKX2*.*5* and *GATA4* showed significantly lower expression levels at day 7, which for *NKX2*.*5*, but not for *GATA4*, recovered at 14 days of differentiation. A similar trend was observed in two (*Myh6 and TNNI3*) out of three (*TNNT2*) tested late stage cardiomyocyte differentiation markers. These delaying of cardiomyocyte differentiation by Bcl-2 KO either suggest that that Bcl-2 is required for the efficient upregulation of cardiac differentiation markers or that Bcl-2 is contributes to increasing/maintaining the number of differentiated cardiomyocytes, which in turn mediates the increased expression of these cardiac differentiation markers.

To further support these observations and show that KO of Bcl-2 does not impair cardiomyocyte differentiation from hiPSC cultures were immuonstained for cardiac troponin T (cTnT) after 14 days of differentiation. In both control and KO conditions, we were able to detect areas of cTNT staining (Figure 2C) supporting the observation that Bcl-2 does not completely abrogate the differentiation of cardiomyocytes from hiPSC.

Bcl-2 KO does not trigger cell death induction or changes in autophagic flux in hiPSC-derived cardiomyocytes One explanation for the lower expression of cardiac progenitor and late cardiac differentiation markers in the Bcl-2 KO conditions could be that Bcl-2 KO results in increased cell death, which leads to a lower number of differentiated cells and an impact on population measures of differentiation. This is indeed plausible as Bcl-2 is known to play important functions in apoptosis and autophagy, two pathways which may control cell numbers.

Bcl-2 is a key anti-apoptotic protein known to bind to and inhibit pro-apoptotic Bax and Bak, crucial for the execution of apoptosis. Therefore, KO of Bcl-2 may indeed lead to increased apoptotic cell death. In order to address this, we performed immunoblot experiments and stained for poly ADP-ribose polymerase (PARP), which is cleaved by active caspase 3, an effector of apoptosis downstream of Bax/Bak activation. No difference in the ratio of the cleaved to total PARP was observed between control and Bcl-2 KO lines indicating no increase in caspase 3 activity (Figure 3).

**Figure 3:**
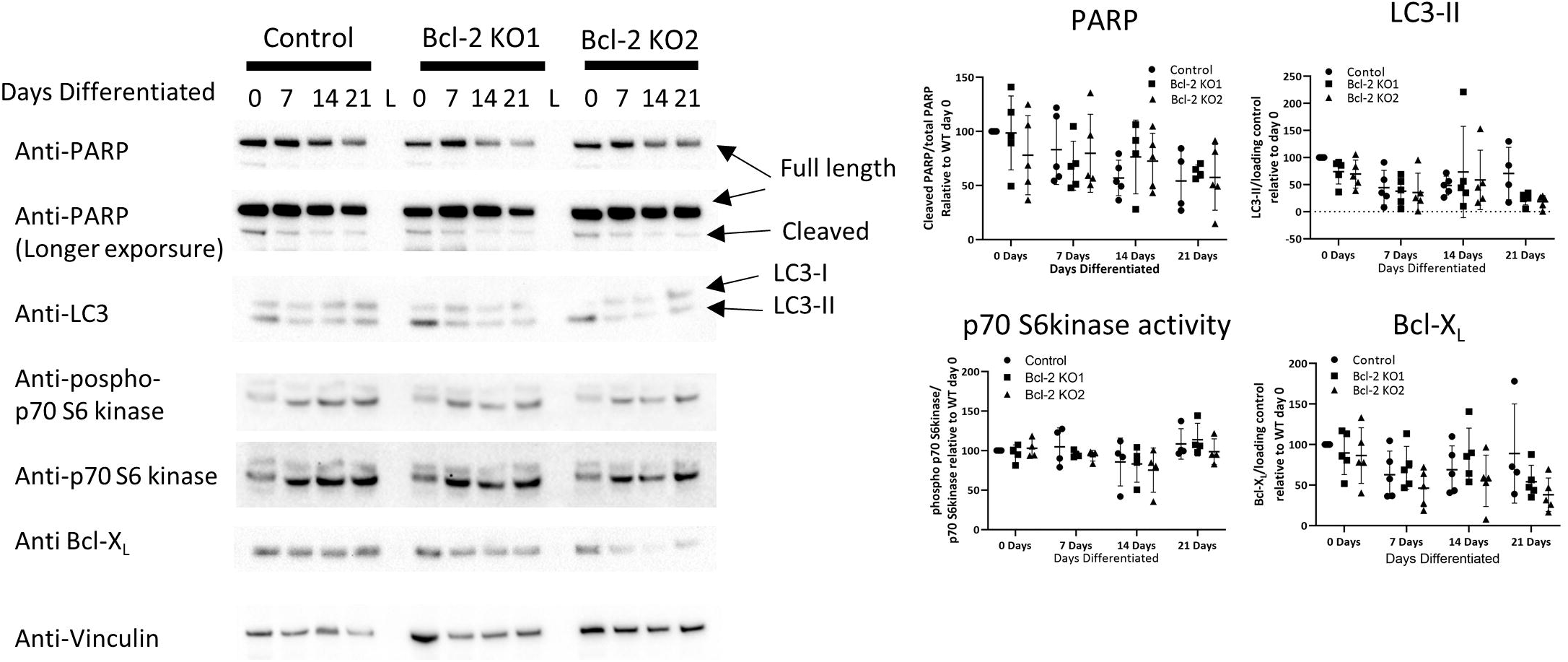
KO of Bcl-2 does not induce apoptosis or changes in autophagic flux. **Left**, immunoblot analysis for the indicated proteins in lysates prepared from cardiomyocytes derived from control or Bcl-2 KO clones differentiated for 0, 7, 14 or 21 days. **Right**, quantification of the performed immunoblot experiments. For PARP cleavage, the expression levels of cleaved PARP were normalised to the total PARP levels. For p70 S6 kinase activity, the phosphorylated p70 S6 kinase levels were divided by the total p70 S6kinase levels. All other protein expression levels are plotted normalized to their corresponding loading control (vinculin). All protein levels are shown relative to the levels for control day 0. All differentiations were performed at least 4 times (n≥4) the average ±SD is indicated in the figures. For comparing day 0 values, a non-parametric Kruskall-Wallis tests with Dunn/s multiple comparison was performed to compare to the normalized control value of control day 0. For all other days, one-way ANOVA tests with Tukey-post test for multiple comparison were performed as the assumptions of normality were met in these conditions.

Bcl-2 also plays important roles in the regulation of autophagy. By binding to Beclin 1 for instance, Bcl-2 impairs the induction of autophagosome formation thus inhibiting autophagic flux. In order to measure autophagy induction, we monitored the levels of lipidated microtubule-associated protein light chain 3 (LC3-II). No difference in LC3-II levels could be observed between control and Bcl-2 KO clones (Figure 3). In addition, we also monitored mammalian target of rapamycin (mTOR) activity by determining the phosphorylation state/activity of p70 S6 kinase (figure 3), a downstream target of mTOR. Similarly to the LC3-II results the phosphorylation state of p70 S6 kinase was unaltered confirming that Bcl-2 KO did not alter basal autophagic flux induction.

Finally, we validated whether the Bcl-2 KO clones upregulated the closely related anti-apoptotic Bcl-X_L_ protein to compensate for loss of Bcl-2. However, Bcl-X_L_ levels were not significantly altered in the Bcl-2 KO clones compared to the control and even seemed to decrease in all conditions rather than increase during differentiation (Figure 3).

### Bcl-2 KO reduces the area of beating cells and inhibits the amplitude of spontaneous Ca^2+^ transients in hiPSC-derived cardiomyocytes

As the induction of cardiomyocyte differentiation was delayed upon KO of Bcl-2 and no differences were observed in apoptosis and autophagy induction, we next set out to examine whether functional measures were similarly affected in Bcl-2 KO hiPSC-derived cardiomyocytes. To these ends, we investigated spontaneous Ca^2+^ transients, which are responsible for the spontaneous contractions in WT and Bcl-2 KO cardiomyocytes, using live cell Ca^2+^ imaging. Ca^2+^ imaging experiments were performed in cardiomyocytes differentiated for 9 or 11 days. At these time points it was feasible to detect spontaneously contracting areas in both control and Bcl-2 KO cardiomyocytes. Fluo-4 loading allowed the measuring of the surface area of beating cells by monitoring spontaneous release. This is visualised in Figure 4A as maximum intensity Z projections of the first 30 sec (300 images) of a control and the Bcl-2 KO conditions. Quantification of the area of spontaneously active cardiomyocytes showed that in Bcl-2 KO cells this was severely decreased (Figure 4B). Next, we focused on the properties of the Ca^2+^ release. To assess the dynamics of Ca^2+^ transients over the image, pseudo-line scans of the largest area of spontaneously active cells were performed that spanned the duration of the experiment (Figure 4C). These analyses were performed using the Fiji software by “reslicing” the stack over a vertical line in the image (the arrows in Figure 4A indicate where these lines were drawn). From this line stack, fluorescence intensity measurements were made and plotted as normalised fluorescence (F/F0). This was performed either for the entire slice area (black vertical bar Figure 4C) or was focussed on the area with the largest response (red bar, Figure 4C). A representative image of such a line stack for control and Bcl-2 KO clones can be seen in figure 4C. The intensity plot of the corresponding line stacks is shown in figure 4D. This analysis showed that compared to the control, Bcl-2 KO did not alter the frequency of the spontaneous Ca^2+^ release events (Figure 4E). However, the amplitude (Figure 4F) and area under the curve (AUC; Figure 4G) were significantly lower in Bcl-2 KO cardiomyocytes compared to the control cardiomyocytes, when quantifying the entire line stack. From figure 4A, B and the line stack, it is clear that a larger area of control cardiomyocytes exhibit spontaneous activity compared to the Bcl-2 KO clones. We therefore also performed the same quantification but restricted to the major sites of response (red line Figure 4C) and saw that both amplitude and AUC were no longer significantly different between control and Bcl-2 KO. These results confirm that KO of Bcl-2 does not impair but rather delays cardiomyocyte differentiation from hiPSC. These data highlight the advantage of performing imaging experiments where cell populations of differing maturation can be discretely analysed.

**Figure 4:**
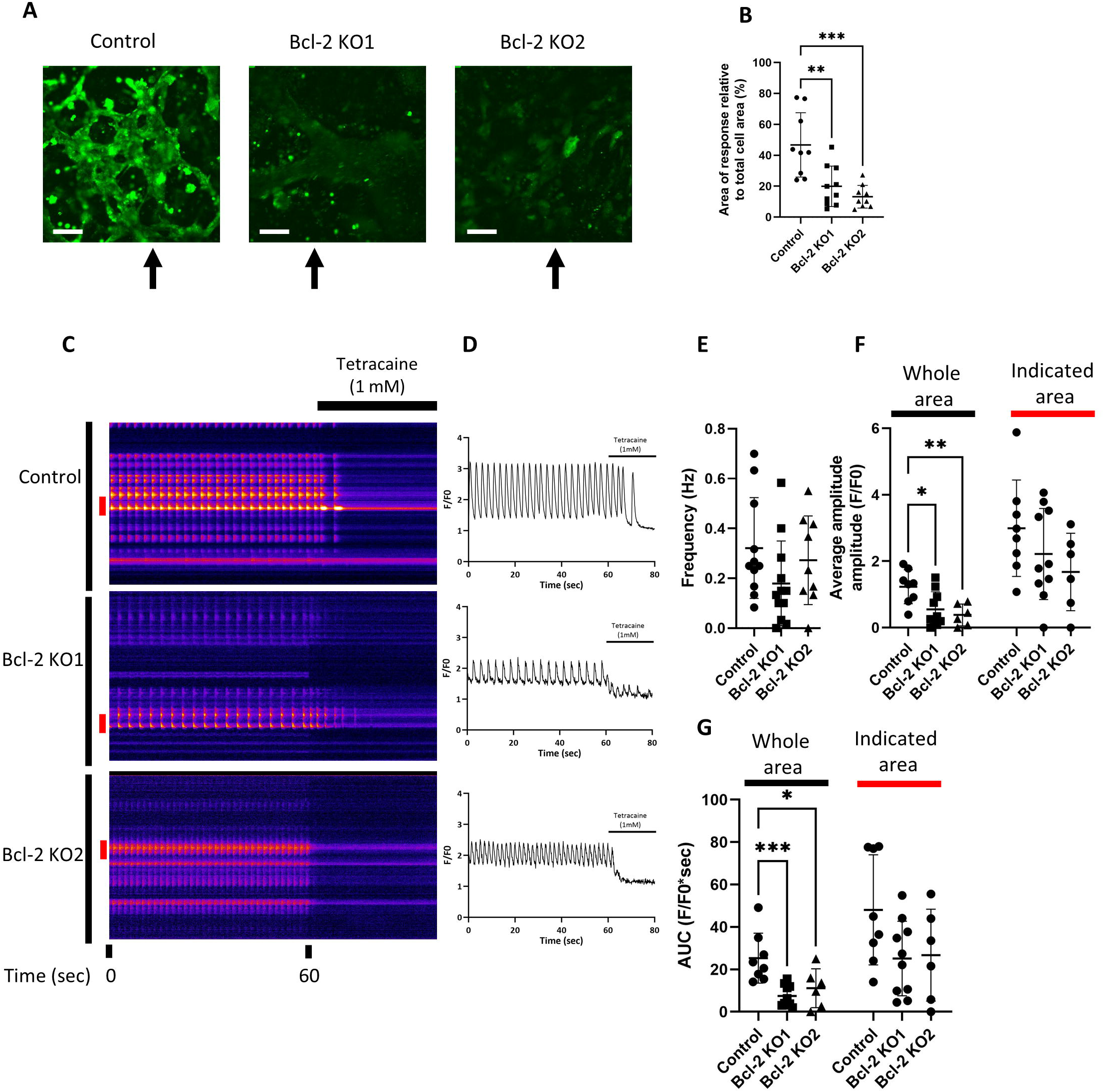
KO of Bcl-2 reduces the area of spontaneous activity as well as the spontaneous Ca2+ release. Spontaneous intracellular Ca^2+^ measurements in cardiomyocytes differentiated for 9 or 11 days using Fluo-4. **A)** Maximum intensity projections of the first 30 s (300 images) of a representative Ca^2+^ measurement for control and Bcl-2 KO conditions. The white scale bar corresponds to 100 µM. The arrows indicate where the line was drawn for “reslicing” the stack in C. **B)** Quantification of the area of spontaneous activity derived from the maximum intensity plots relative to the total area covered by cells (%). Each data point represents the size of the area of spontaneous activity of a single Ca^2+^ imaging experiment. The average ± SD are indicated on the graphs (n≥9 differentiations?). **C)** Visual representation of the experiment showing a time laps of the pseudo-line scans obtained after re-slicing the image stack for control and Bcl-2 KO conditions. After 60 s, tetracaine (1mM) was added to block RyR activity and determine the baseline **D)** Intensity profiles obtained by quantifying the entire area (indicated by the black bar left of the pseudo-line scans in **C)** plotted as F/F0. F0 was determined after tetracaine addition. Quantification of the spontaneous activity was obtained by determining the frequency **E**, amplitude **F** and area under the curve G. For amplitude and frequency an additional quantification was performed focussing only on the areas of most intense responses (indicated by the red bar left of the line scans in **C)**. All experiments were performed at least 6 times (n≥ 6). The averages ± SD are represented on the graphs. For statistical analysis one-way ANOVA tests with Tukey-post test for multiple comparison were performed. * indicates p<0.05, ** indicates p<0.01, *** indicate p<0.001

### Bcl-2 KO impairs the expression of the cardiac Ca^2+^ toolkit

Our findings suggest that Bcl-2 KO delays differentiation of hiPSC to cardiomyocytes and as a consequence spontaneous Ca^2+^ transients and contraction are inhibited. However, where Bcl-2 KO cells do differentiate they show similar functional activity compared to controls. This suggests Bcl-2 does not block cardiomyocyte differentiation but rather intervenes in deciding whether or not hiPSC differentiate to cardiomyocytes. Given the association between the expression of specific cardiac Ca^2+^ handling machinery involved in excitation contraction coupling with cardiomyocyte maturation we next investigated the underlying mechanisms for this reduced Ca^2+^ signalling capacity. To this end, immunoblot assays were performed to evaluate the expression levels of a number of proteins involved in cardiac function and Ca^2+^ handling (Figure 5). From these immunoblots it is clear that the expression levels of RyR2, the sodium Ca^2+^ exchanger (NCX), the cardiac specific sarco/endoplasmic reticulum ATPase 2A (SERCA2a) isoform and cardiac cTnT were dramatically lower in Bcl-2 KO cardiomyocytes compared to control cardiomyocytes. Inositol 1,4,5 Trisphosphate receptor type 2 (IP_3_R2) levels showed a similar decline in control and Bcl-2 KO conditions, indicating the decrease is specific for cardiac specific markers. These results suggest that when examining the entire cell population in the Bcl-2 KO cultures, only a minority is fully differentiated to cardiomyocytes. By focussing on the spontaneously active regions in the Ca^2+^ imaging experiments, we most likely underestimate the effects of Bcl-2 KO on cardiomyocyte Ca^2+^ handling.

**Figure 5:**
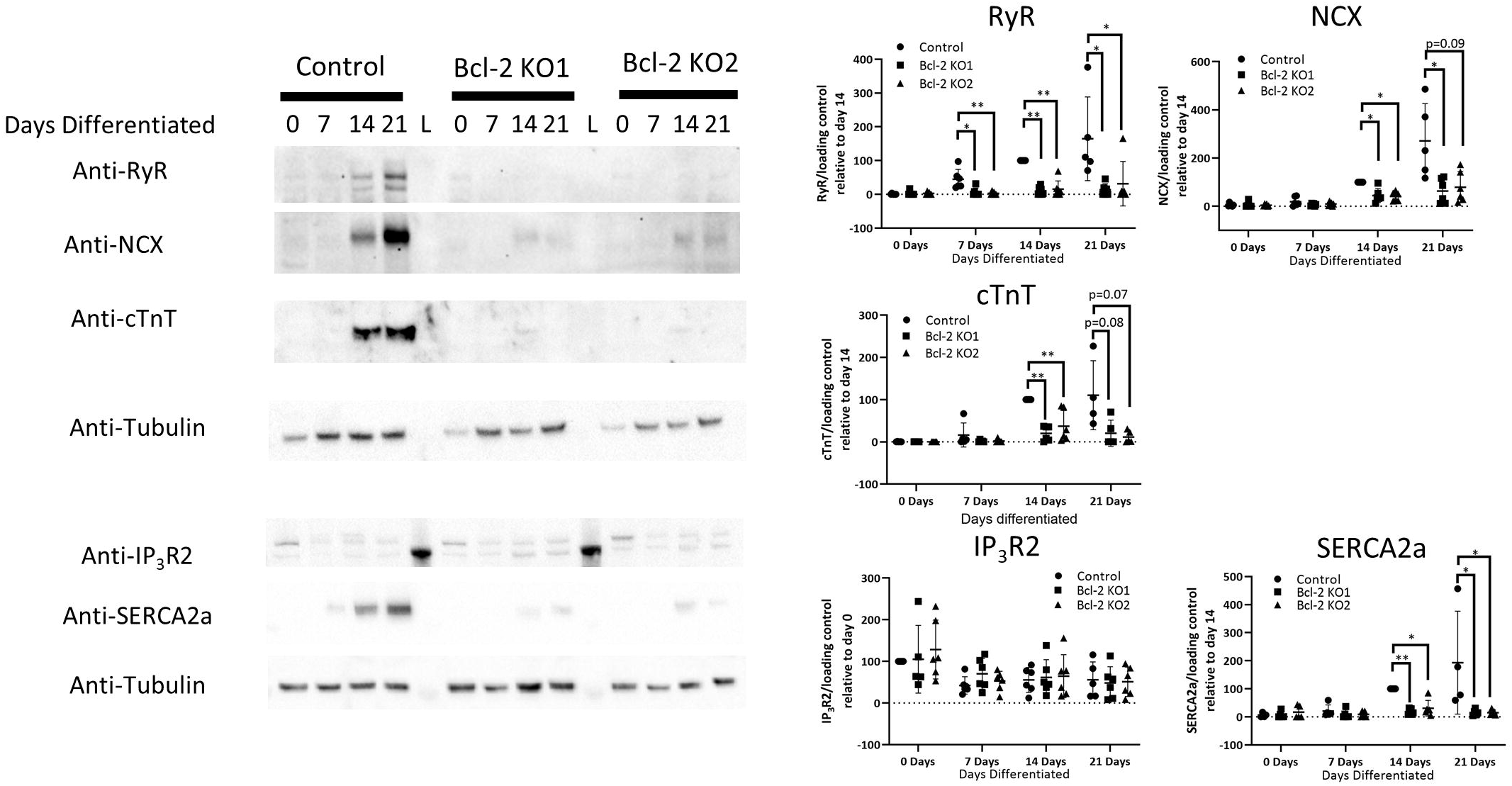
KO off Bcl-2 does impairs the expression of the cardiac Ca^2+^ toolkit. Left, Immunoblot analysis of the indiacted proteins in lysates prepared from cardiomyocytes derived from control or Bcl-2 KO clones differentiated for 0, 7, 14 or 21 days. Right, quantification of the performed immunoblot experiments. All protein expression levels are normalized towards their corresponding loading control (tubulin). On each blot all values were normalized to a control condition where expression levels were easily detected. For RyR, SERCA2a, cTNT, NCX this was control day 14 as for IP_3_R2 this was control day 0. Each data point represents and independent differentiation which were performed at least 4 times (n≥4). The averages ± SD are represented on the graphs. As the assumption for normal distribution were not met for all conditions, non-parametric Kruskall-Wallis tests with Dunn’s multiple comparison were performed * indicates p<0.05 and ** indicates p<0.01.

### Bcl-2 KO impairs early c-Myc upregulation thereby resulting in reduced functional cardiomyocyte growth

KO of Bcl-2 did not alter the levels of apoptosis or basal autophagic flux. Besides participating in pathways involved in cell removal, Bcl-2 is also involved in the regulation of cellular proliferation differentiation and growth. An interesting link between Bcl-2 and c-Myc has been described whereby Bcl-2 would stabilize c-Myc thus increasing its expression and activity (13). To explore this possibility in hiPSC-derived cardiomyocytes, we determined c-Myc levels between control and Bcl-2 KO lines (Figure 6A). Interestingly, on differentiation day 7, c-Myc is upregulated in control cells which was not the case in both Bcl-2 KO cell lines compared to the control condition. A similar non-significant trend was observed at day 14 whereas at day 21 c-Myc expression levels recovered to control levels. This observation indeed suggests that loss of Bcl-2 results in decreased c-Myc expression early during cardiomyocyte differentiation from hiPSC.

**Figure 6:**
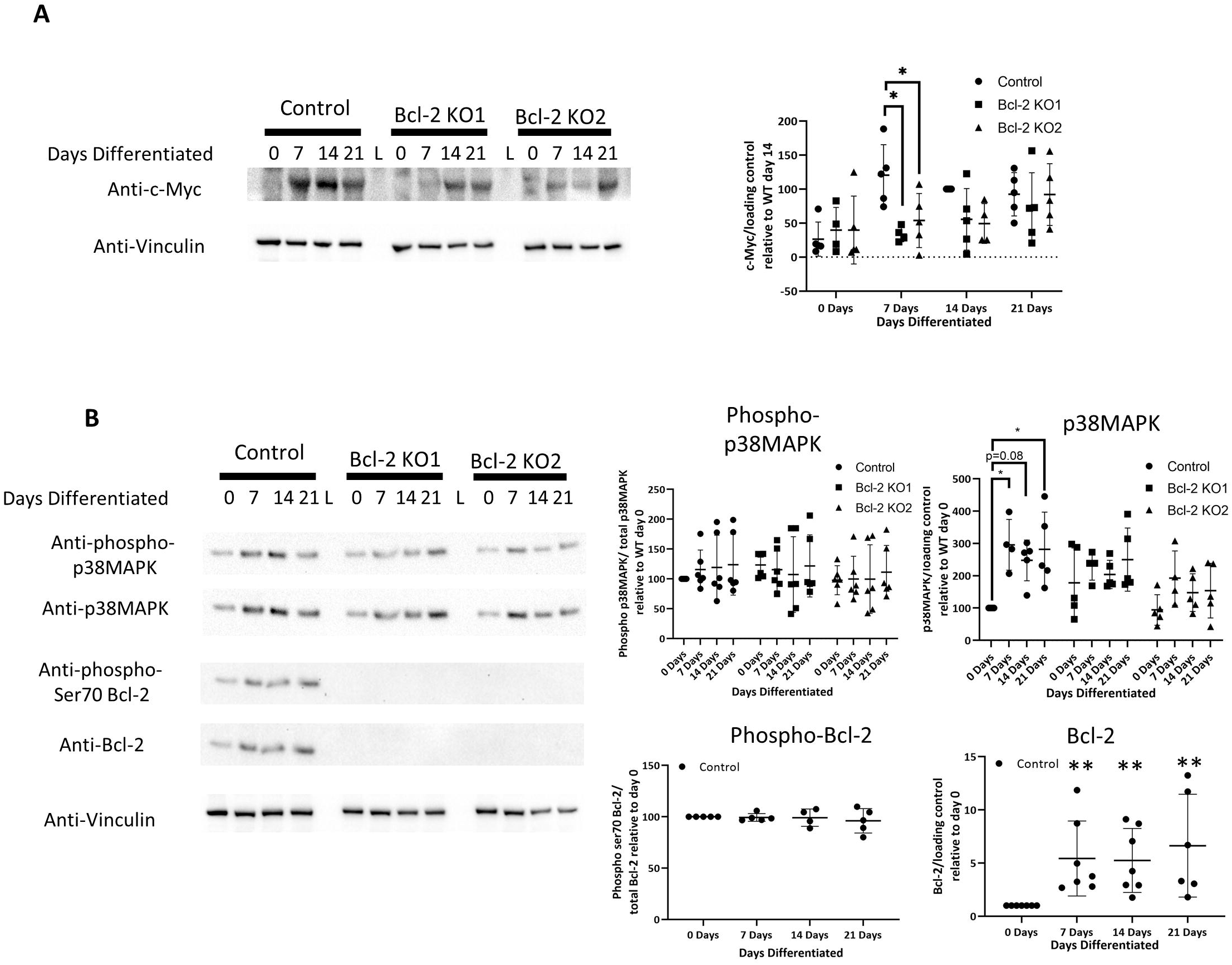
Bcl-2 KO impairs upregulation of c-Myc. **A**,**B)** Left, immunoblot analysis of cardiomyocytes derived from control or Bcl-2 KO clones differentiated for 0, 7, 14 or 21 days, stained for the indicated proteins. **Right**, quantification of the performed immunoblot experiments. For quantifying phosphorylated p38MAPK or phosphorylated Bcl-2, the protein levels detected by the phospho-antibody were normalized to the total amount of p38MAPK or Bcl-2. For the other quantifications, protein levels were normalized to their respective loading control (vinculin) and shown relative to a control condition were the protein was easily detected. For Bcl-2 and p38MAPK, this was control day 0 whereas for c-Myc, this was control day 14. All differentiations (represented by a data point) were performed at least times (n≥4). The averages ± SD are represented on the graphs. In the conditions normalized to, the assumptions for normal distribution were not met. Therefore, when comparing these conditions Kruskall-Wallis tests with Dunn’s multiple comparison were performed. All other conditions were normally distributed and allowed the use of one-way ANOVA tests with Tukey-post test for multiple comparison were performed. * indicates p<0.05, ** indicates p<0.01.

It has been shown that phosphorylation of Bcl-2 by MAPK, a kinase heavily involved in regulating cell proliferation, leads to the nuclear translocation of Bcl-2 where it provides pro-survival signalling and chemoresistance (12). This effect of Bcl-2 could at least be in part due to its stabilizing effects on c-Myc in the nucleus thus promoting c-Myc signalling (13). To validate the involvement of this pathway here, we determined levels of the active phosphorylated form of p38MAPK and of phosphorylated Bcl-2 by immunoblot analysis (Figure 6B). Levels of phosphorylated active p38MAPK normalised to total p38MAPK levels were not altered between non-differentiated control and Bcl-2 KO cells. However, during differentiation p38MAPK expression levels significantly increased in control cardiomyocytes signifying an increase in absolute levels of active p38MAPK. No such increase was detected during the differentiation of the Bcl-2 KO cells. Similarly, Bcl-2 phosphorylated at serine 70, which is a known target of MAPK pathways and which facilitate its nuclear translocation (12), could be detected throughout the differentiation of WT hiPSC (Figure 6B). Interestingly, in the cardiomyocytes derived from WT hiPSC, Bcl-2 levels were upregulated from day 7 onward coinciding with the increased levels/activity of p38MAPK and c-Myc, highlighting the importance of Bcl-2 for cardiac differentiation

These results suggest that Bcl-2 KO results in reduced expression of c-Myc early in cardiomyocyte differentiation. As c-Myc is an important regulator of cardiomyocyte differentiation this may thus account at least in part for the delays in differentiation observed in the Bcl-2 KO hiPSC derived cardiomyocytes.

## Discussion

In this study, we show that Bcl-2 plays an important part in the early phases of cardiomyocyte differentiation from hiPSC. KO of Bcl-2 did not induce apoptosis or alter autophagic flux induction but instead resulted in delayed differentiation of cardiomyocytes from hiPSC. Our result point towards a role for Bcl-2 in regulating early c-Myc stability, a transcription factor known to be required for cardiac differentiation and growth (16).

It is known that Bcl-2 plays important roles during differentiation of neurons and cardiomyocytes however the pathways involved remain to be fully elucidated (5-8). In this study we aimed to address this further and build on known functions of Bcl-2 in cardiomyocyte differentiation. To this end, a CRISPR/Cas9-mediated knock-out of Bcl-2 was performed and validated in hiPSC (Figure 1). The CRSIPR/Cas9 assay that was employed in this study was intended to introduce specific point mutations via homologues recombination in the *BCL2* gene resulting in the introduction of premature stop codons to prevent protein translation. Insertion of the selection cassette carrying the mutations was successful at the intended locus. However, after removal of the selection cassette none of the tested clones showed the introduction of the intended mutations. Rather, the majority of the clones showed a 4 base pair deletion at the gRNA cut site resulting in a frame shift and insertion of three stop codons, which effectively resulted in knocking-out Bcl-2. The lack of protein was confirmed by immunoblot, which showed a complete KO of Bcl-2 compared to the control line indicating that these clones were suitable for use in this study.

After validating the KO of Bcl-2, the effects of lack of Bcl-2 on cardiomyocyte differentiation were addressed. In our differentiation of hiPSC to cardiomyocytes, a delay in the expression profile of four out of five tested early and late markers associated with cardiomyocyte differentiation was observed comparing control and Bcl-2 KO (Figure 2). This suggests that KO of Bcl-2 delays the differentiation of cardiomyocytes from hiPSC. This was further confirmed by measuring spontaneous intracellular Ca^2+^ release, events as a measure of cardiomyocyte differentiation and maturation, in addition to measuring the size of areas spontaneous activity (Figure 4). From these experiments it was clear that control cardiomyocytes showed larger spontaneous Ca^2+^ release over larger areas compared to the Bcl-2 KO conditions. However, when focused exclusively on the major site of activity no significant functional differences between control and Bcl-2 KO conditions could be detected. These results suggest that KO of Bcl-2 does not impair cardiomyocyte differentiation but rather influences the number of cells that successfully differentiate. This also fits well with the observation that Bcl-2 KO delays differentiation. This then limits the number of functional cardiomyocytes during the early stages of differentiation. A consideration here is that we are likely underestimating the effects of Bcl-2 KO on Ca^2+^ signalling. In these experiments, the decision to measure an area was made based on the presence of visual spontaneous activity. This was much more pronounced in the control cardiomyocytes compared to the Bcl-2 KO. As such in the Bcl-2 KO we are likely sampling a sub population of the cells that actually did differentiate properly and as such also does not show large defects when specifically quantified.

The latter is further confirmed by the lack of expression of critical members of the cardiac Ca^2+^ toolkit in Bcl-2 KO cardiomyocytes (Figure 5). These immunoblots monitor the average of the entire differentiating population. Based on the largely absent RyR, cTnT, SERCAa1 and NCX in the Bcl-2 KO condition one would not expect any spontaneous activity to be possible in the Bcl-2 KO cardiomyocyte. Nevertheless, a subpopulation still was able to produce spontaneous Ca^2+^ release and contractions (Figure 4). Furthermore, Bcl-2 KO did not result in excessive induction of apoptosis or changes in autophagic flux (Figure 3). Taken together these results indicate that Bcl-2 KO delays differentiation limiting the number of functional cardiomyocytes without affecting death of cardiomyocytes. This might be a more general function of Bcl-2, not only restricted to cardiomyocytes, as Bcl-2 KO mice have severe growth retardation which may be attributed to a general delay in differentiation when Bcl-2 is not present (20).

During cardiomyocyte differentiation, both p38MAPK and c-Myc activity play important roles in differentiation and growth. p38MAPK has been shown to act as a switch deciding between proliferation and differentiation of cardiomyocytes (14, 15, 21). c-Myc expression in cardiomyocytes on the other hand has been shown to be involved in cardiac growth (16). Bcl-2 interacts with nuclear c-Myc thereby regulating its transcriptional activity and stability (13). Bcl-2 translocation to the nucleus is dependent on its phosphorylation status (12). p38MAPK phosphorylates Bcl-2 and as such may facilitate its translocation towards the nucleus (11). Other MAPK are also known to phosphorylate Bcl-2. ERK1/2 for instance phosphorylates Bcl-2 at Ser70 thereby impacting its anti-apoptotic functions (22) and its functional cooperation with c-Myc (23), suggesting this is an important mechanism for several MAPK in the regulation of Bcl-2. Our results show that before and during differentiation in both control and Bcl-2 KO conditions, p38MAPK is active and able to phosphorylate its targets (Figure 6B). In control conditions, Bcl-2 is phosphorylated and upregulated as has been previously described (7). Together with Bcl-2, c-Myc levels are significantly higher at day 7 in control compared to KO conditions (Figure 6A, B). Prolonged differentiation results in recovery of c-Myc levels towards control levels in both KO lines. c-Myc regulates the expression of many genes (24). It is therefore logical that reducing c-Myc levels would result in a reduction of c-Myc driven transcription and potentially reduced cardiac differentiation. Recently it was shown that transcriptional output of c-Myc in the heart, following upregulation of c-Myc, is largely dependent on the presence of transcriptional cofactors determining which genes will be activated (25). Furthermore, the abundance of these cofactors was shown to be a determining/rate limiting factor as to which genes are regulated by c-Myc (25). In our hands we see a reduction in the abundance of c-Myc upon Bcl-2 KO thus suggesting that lack of cofactors would not be an issue in this setting. However, in the context of cardiac differentiation of the Bcl-2 KO hiPSC, these cardiac specific cofactors are expected to direct the limited amount of c-Myc towards genes necessary for the differentiation, thereby allowing this process to occur at a slow/delayed rate compared to the control condition. This may at least in part explain the delay in differentiation observed upon Bcl-2 KO.

In summary, our results indicate that Bcl-2 is required for maintaining the temporal trajectory of early cardiomyocyte differentiation by regulating c-Myc expression. Loss of Bcl-2 thus results in delayed differentiation of cardiomyocytes from hiPSC resulting in less efficient accumulation of functional cardiomyocytes.

## Acknowledgments

This work was supported by the Research Foundation-Flanders (FWO) “krediet aan navorsers” (grant number: 1508319N). TV is a recipient of a senior post-doctoral grant of the FWO (grant number: 12ZG121N)

## References

1. Brunelle JK, Letai A. Control of mitochondrial apoptosis by the Bcl-2 family. J Cell Sci. 2009;122(4):437–41.

2. Rong YP, Aromolaran AS, Bultynck G, Zhong F, Li X, McColl K, et al. Targeting Bcl-2-IP3 receptor interaction to reverse Bcl-2’s inhibition of apoptotic calcium signals. Mol Cell. 2008;31(2):255–65.

3. Vervliet T, Parys JB, Bultynck G. Bcl-2 proteins and calcium signaling: complexity beneath the surface. Oncogene. 2016;35(39):5079–92.

4. Pattingre S, Tassa A, Qu X, Garuti R, Liang XH, Mizushima N, et al. Bcl-2 antiapoptotic proteins inhibit Beclin 1-dependent autophagy. Cell. 2005;122(6):927–39.

5. Wei L, Cui L, Snider BJ, Rivkin M, Yu SS, Lee CS, et al. Transplantation of embryonic stem cells overexpressing Bcl-2 promotes functional recovery after transient cerebral ischemia. Neurobiol Dis. 2005;19(1-2):183–93.

6. Liang Y, Mirnics ZK, Yan C, Nylander KD, Schor NF. Bcl-2 mediates induction of neural differentiation. Oncogene. 2003;22(35):5515–8.

7. Kobayashi S, Lackey T, Huang Y, Bisping E, Pu WT, Boxer LM, et al. Transcription factor gata4 regulates cardiac BCL2 gene expression in vitro and in vivo. FASEB J. 2006;20(6):800–2.

8. Limana F, Urbanek K, Chimenti S, Quaini F, Leri A, Kajstura J, et al. bcl-2 overexpression promotes myocyte proliferation. Proc Natl Acad Sci U S A. 2002;99(9):6257–62.

9. Maundrell K, Antonsson B, Magnenat E, Camps M, Muda M, Chabert C, et al. Bcl-2 undergoes phosphorylation by c-Jun N-terminal kinase/stress-activated protein kinases in the presence of the constitutively active GTP-binding protein Rac1. J Biol Chem. 1997;272(40):25238–42.

10. Blagosklonny MV. Unwinding the loop of Bcl-2 phosphorylation. Leukemia. 2001;15(6):869–74.

11. De Chiara G, Marcocci ME, Torcia M, Lucibello M, Rosini P, Bonini P, et al. Bcl-2 Phosphorylation by p38 MAPK: identification of target sites and biologic consequences. J Biol Chem. 2006;281(30):21353–61.

12. Zhou M, Zhang Q, Zhao J, Liao M, Wen S, Yang M. Phosphorylation of Bcl-2 plays an important role in glycochenodeoxycholate-induced survival and chemoresistance in HCC. Oncol Rep. 2017;38(3):1742–50.

13. Jin Z, May WS, Gao F, Flagg T, Deng X. Bcl2 suppresses DNA repair by enhancing c-Myc transcriptional activity. J Biol Chem. 2006;281(20):14446–56.

14. Aouadi M, Bost F, Caron L, Laurent K, Le Marchand Brustel Y, Binetruy B. p38 mitogen-activated protein kinase activity commits embryonic stem cells to either neurogenesis or cardiomyogenesis. Stem Cells. 2006;24(5):1399–406.

15. Engel FB, Schebesta M, Duong MT, Lu G, Ren S, Madwed JB, et al. p38 MAP kinase inhibition enables proliferation of adult mammalian cardiomyocytes. Genes Dev. 2005;19(10):1175–87.

16. Jackson T, Allard MF, Sreenan CM, Doss LK, Bishop SP, Swain JL. The c-myc proto-oncogene regulates cardiac development in transgenic mice. Mol Cell Biol. 1990;10(7):3709–16.

17. Chu VT, Weber T, Wefers B, Wurst W, Sander S, Rajewsky K, et al. Increasing the efficiency of homology-directed repair for CRISPR-Cas9-induced precise gene editing in mammalian cells. Nat Biotechnol. 2015;33(5):543–8.

18. Vervliet T, Decrock E, Molgo J, Sorrentino V, Missiaen L, Leybaert L, et al. Bcl-2 binds to and inhibits ryanodine receptors. J Cell Sci. 2014;127(Pt 12):2782–92.

19. Rueden CT, Schindelin J, Hiner MC, DeZonia BE, Walter AE, Arena ET, et al. ImageJ2: ImageJ for the next generation of scientific image data. BMC Bioinformatics. 2017;18(1):529.

20. Veis DJ, Sorenson CM, Shutter JR, Korsmeyer SJ. Bcl-2-deficient mice demonstrate fulminant lymphoid apoptosis, polycystic kidneys, and hypopigmented hair. Cell. 1993;75(2):229–40.

21. Romero-Becerra R, Santamans AM, Folgueira C, Sabio G. p38 MAPK Pathway in the Heart: New Insights in Health and Disease. Int J Mol Sci. 2020;21(19).

22. Deng X, Ruvolo P, Carr B, May WS, Jr. Survival function of ERK1/2 as IL-3-activated, staurosporine-resistant Bcl2 kinases. Proc Natl Acad Sci U S A. 2000;97(4):1578–83.

23. Jin Z, Gao F, Flagg T, Deng X. Tobacco-specific nitrosamine 4-(methylnitrosamino)-1-(3-pyridyl)-1-butanone promotes functional cooperation of Bcl2 and c-Myc through phosphorylation in regulating cell survival and proliferation. J Biol Chem. 2004;279(38):40209–19.

24. Patel JH, Loboda AP, Showe MK, Showe LC, McMahon SB. Analysis of genomic targets reveals complex functions of MYC. Nat Rev Cancer. 2004;4(7):562–8.

25. Bywater MJ, Burkhart DL, Straube J, Sabo A, Pendino V, Hudson JE, et al. Reactivation of Myc transcription in the mouse heart unlocks its proliferative capacity. Nat Commun. 2020;11(1):1827.

