## Supplemental information for "Cardiomyocyte differentiation from iPS cells is delayed following knockout of Bcl-2"

**Supplemental materials and methods**

*BCL2* homologues sequence (HAL)

Tgtaaattgccgagaaggggaaaacatcacaggacttctgcgaataccggactgaaaattgtaattca
tctgccgccgccgctgcctttttttttttctcgagctcttgagatctccggttgggattcctgcggatt
gacatttctgtgaagcagaagtctgggaatcgatctggaaatcctcctaattttactccctctcccc
gcgactcctgattcattgggaagtttcaaatcagctataactggagagtgctgaagattgatgggatc
gttgccttatgcatttgttttggttttacaaaaaggaaacttgacagaggatcatgctgtacttaaaa
aatacaagtaagttctctgcacaggaaattggtttaatgtaactttcaatggaaacctttgagatttt
ttacttaaagtgcattcgagtaaatttaatttccaggcagc

*BCL2* homologues sequence (HAR)

tacattctttttagccgtgttacttgtagtgtgtatgccctgctttcactcagtgtgtacagggaaac
gcacctgatttttttacttattagtttgtttttttctttaacctttcagcatcacagaggaagtagactg
atattaacaatacttactaataataacgtgcctcatgaaataaagatccgaaaggaattggaataaaa
atttcctgcatctcatgccaaggggggaaacaccagaatcaagtgttccgcgtgattgaagacaccccc
tcgtccaagaatgcaaagcacatccaataaaatagctggattataactcctcttctttctctgggggc
cgtggggtgggagctggggcgagaggtgccgttggcccccgttgcttttcctctgggaaggatggcgt
aagctgggagaacagggtacgataaccgtgagatagtgtaaa

**Supplemental table 1: List of primers used for genomic DNA screening**

| Gene | Primer direction | Primer sequence (5’ > 3’) |
|---|---|---|
| *BCL2* | forward | GTGCTGAAGATTGATGGGATCG |
| | reverse | CTCAAAGAAGGCCACAATCCTCC |
| *HAL insertion* | forward | GCGCGTCCTGCCTTCATTTATCC |
| | reverse | GACACTTACCGCATTGACAAGCACG |
| *HAR insertion* | forward | AGGCGGGCCATTTACCGTAAG |
| | reverse | GAGGAGAAGATGCCCGGTGC |
| *RFBOX* | forward | AGTCCCAGCCCTCTAATCACAAAG |
| | reverse | GGTGACTTTGGACAGGTGGCTCAG |
| *SLC2A9* | forward | ATTCTCACAACATCCCTGCCAGTGC |
| | reverse | GAAGGTGCAACACAATGACTCTGG |
| *ANKRD28* | forward | CAGACAGTGATCCTTAGGCTTC |
| | reverse | CACAAGGCGGAAATAGTCTGGCACC |
| *NEFL* | forward | GTTCAGGATCTACGGCAATGTG |
| | reverse | TGCAATGTCCAACCAGTCAAGC |
| *SLC23A* | forward | CCTAACTAATACAGCCCTCACTGG |
| | reverse | CCAGGCTCGGGTGAGGGAGTTAC |
| *PTPRT* | forward | AGCATTGCCAACCACTAGCAGAAG |
| | reverse | TGATGCGGATGAAGAGGCTGAGTC |
| *NCKAP5* | forward | GAAGGCAGGGATAGGGCAGGACAAG |
| | reverse | GCAAAGTGAGGAGACGAGATAACC |
| *FRAS1* | forward | AAGCACAGGACCAGACTTGCCAG |
| | reverse | GGCTGGCAGTTTGGAACAGGTGTG |
| *DLG2* | forward | GAGGAAATGGATGGAGAGTGAGG |
| | reverse | AAGCCTTCAGGGAGTGACCATC |
| *PCDH19* | forward | CAGCAATCGACTCCAAAGAACC |
| | reverse | TTCACTGAGCCTAACCACCAAG |

**Supplemental table 2:** List of primers utilized in this study for Quantitative Real-Time PCR.

| Gene | Primer direction | Primer sequence (5' > 3') |
|---|---|---|
| *OCT4* | forward | CGAGCAATTTGCCAAGCTCCTGAA |
| | reverse | GCCGCAGCTTACACATGTTCTTGA |
| *NANOG* | forward | TGGCCGAAGAATAGCAATGGTGTG |
| | reverse | TTCCAGGTCTGGTTGCTCCACATT |
| *BRACH* | forward | ACCCAGTTCATAGCGGTGAC |
| | reverse | AAGCTTTTGCAAATGGATTG |
| *GATA4* | forward | CGACACCCCAATCTCGATATG |
| | reverse | GTTGCACAGATAGTGACCCGT |
| *NKX2.5* | forward | ACCTCAACAGCTCCCTGACTCT |
| | reverse | ATAATCGCCGCCACAAACTCTCC |
| *MYH6* | forward | GCCCTTTGACATTCGCACTG |
| | reverse | CGGGACAAAATCTTGGCTTTGA |
| *TNNI3* | forward | GATGCGGCTAGGGAACCTC |
| | reverse | GCATAAGCGCGGTAGTTGGA |
| *TNNT2* | forward | ACAGAGCGGAAAAGTGGGAAG |
| | reverse | TCGTTGATCCTGTTTCGGAGA |
| *GAPDH* | forward | TCAAGAAGGTGGTGAAGCAGG |
| | reverse | ACCAGGAAATGAGCTTGACAAA |
| *RPL13a* | forward | CCTGGAGGAGAAGAGGAAAGAGA |
| | reverse | TTGAGGACCTCTGTGTATTTGTCAA |
